# geneSTRUCTURE: A Modern Platform for Visualization of Gene Structures

**DOI:** 10.64898/2026.03.05.709980

**Authors:** Shumpei Hashimoto, Kentaro Yamada, Takeshi Izawa

## Abstract

Even in the era of genomics and pan-genomics, the gene remains the fundamental unit of heredity. Accurate visualization of gene structures provides essential insights into the organization, regulation, and evolution of each gene. Representing elements such as exons, introns, untranslated regions, and functional domains in a clear and interpretable format is particularly important for analyzing complex gene architectures and for communicating results effectively. Despite the availability of several visualization tools, many are limited in their ability to incorporate user-defined annotations or to support interactive and customizable figure generation, which reduces their utility in modern analytical workflows. To address these limitations, we developed geneSTRUCTURE, both command-line-interface and web-based application designed to provide flexible and user-friendly visualization of gene structures based on widely used annotation formats, GFF3 and GTF. In addition to visualizing core gene components, the platform allows users to overlay supplemental annotations, including mutation sites and protein domains, and to adjust layout features in real time. By combining a modern interface with annotation flexibility and high-resolution output, geneSTRUCTURE offers a robust solution for gene-level visualization.

## Introduction

The advent of long read sequencing technology has fundamentally reshaped the landscape of genomic research (Jung et al., 2019; Rhoads and Au, 2015; Wang et al., 2025, 2021). The technology precipitated a paradigm shift from single-reference analysis to pan-genomic studies (Bayer et al., 2020; Schreiber et al., 2024). By assembling and comparing multiple genomes, these studies have greatly improved our understanding of genetic variation, structural differences, and evolutional processes (Bayer et al., 2020; Schreiber et al., 2024). While pan-genomics has changed genomic research fundamentally, genes themselves remain the basic units of heredity (Morgan et al., 1915). No matter how complete a genome assembly is, it is individual genes and their structure, function, and regulation that ultimately determine the biological traits. Therefore, in our current era of comparative genomics and large-scale variation studies, understanding genes at the individual level remains crucial.

The visualization of gene features such as composition and position of exons and introns for genes offers visual presentation for biologists to integrate annotation, and also helps them to produce high-quality figures for publication. Accordingly, several web applications or software including FancyGene (Rambaldi and Ciccarelli, 2009), GECA (Fawal et al., 2012), FeatureStack (Frech et al., 2012), GSDraw (Wang et al., 2013), GPViz (Snajder et al., 2013), GenePainter (Hammesfahr et al., 2013), and GSDS 2.0 have been developed so far. One of the most widely used platforms is GSDS (Gene Structure Display Server), which has served the scientific community since 2007 and received a major update with version 2.0 in 2015 (Hu et al., 2015). GSDS 2.0 added support for various annotation formats, including GFF and BED, and provided a visual editor for basic customization. However, GSDS and similar older tools have limitations that increasingly restrict their usefulness as “modern” applications. The GSDS interface has not received major updates since 2015, and its frontend technologies are now outdated. Customization options are limited, especially when researchers want to add extra annotations like sequence variants, deletions, or conserved protein domains to gene models. Additionally, many existing tools lack the interactive features that modern web technologies can provide, which creates steep learning curves and reduces user efficiency.

Here, we developed geneSTRUCTURE, a powerful and highly customizable command-line tool for gene structure visualization. Unlike existing tools, geneSTRUCTURE supports multiple annotation layers, including mutations (insertions, deletions, and SNPs) and multiple domains, with extensive options for fine-tuning visual output. We further released geneSTRUCTURE+, a web-based version featuring a sophisticated user interface (UI) that allows users to edit and customize visualizations interactively in real time. Both tools provide an improved user experience (UX) that meets modern web application standards, enabling researchers to generate publication-ready figures with minimal effort.

## Materials and Methods

### Software architecture and deployment

geneSTRUCTURE was developed as a command-line tool in Python 3.12.6 to parse genome annotation files and generate gene structure figures. geneSTRUCTURE+ provides a web-based interface to geneSTRUCTURE by leveraging FastAPI (v0.115.0) to serve its core functions as a RESTful API. The frontend was implemented in TypeScript using (v14.2.13) with React (v18.3.1), and Mantine was adopted as the UI component library to provide a consistent interface. Dependency management was handled via npm for the frontend and pip for the Python backend, enabling reproducible installation across platforms. The web application was deployed on Vercel and is publicly accessible at https://gene-structure.vercel.app/. The source code and documentation are publicly available on GitHub: the command-line tool at https://github.com/hashimotoshumpei/geneSTRUCTURE and the web application at https://github.com/bvv-1/gene-structure.

## Results and Discussion

geneSTRUCTURE is a tool designed to generate high-quality visualizations of gene structures with user-friendly and flexible customization. It is provided in two formats: a command-line interface (CLI) version, simply referred to as geneSTRUCTURE, and a web-based application, geneSTRUCTURE+ (Fig. 1). Although both versions share essentially the same core functionality, the web-based version offers superior usability in terms of user interface (UI) and user experience (UX). In the CLI version, users first specify an annotation file and provide a CSV file containing transcript IDs along with optional annotations, including deletions, insertions, SNPs, and protein domains specified by amino acid coordinates (Fig. 1B). Annotation files can be provided in either Generic Feature Format Version 3 (GFF3) or Gene Transfer Format (GTF) format. This version, which is implemented in Python, is recommended to use when handling large annotation files exceeding (>1 GB) or in environments where web servers are unavailable (Fig. 1A). Upon execution of the command, the gene structure of any specified transcript is rendered automatically (Fig. 1B). In contrast, the web-based version, geneSTRUCTURE+, is implemented using React and provides an interactive interface with enhanced operability (Fig. 1C). Visualization results are updated in real time in response to parameter changes, enabling highly interactive editing and delivering an improved user experience (Fig. 1D). In both versions, visual attributes of gene components (i.e. exons, introns, and untranslated regions (UTRs)) can be flexibly customized, and the resulting figures can be exported in SVG or PNG formats.

**Fig 1.**
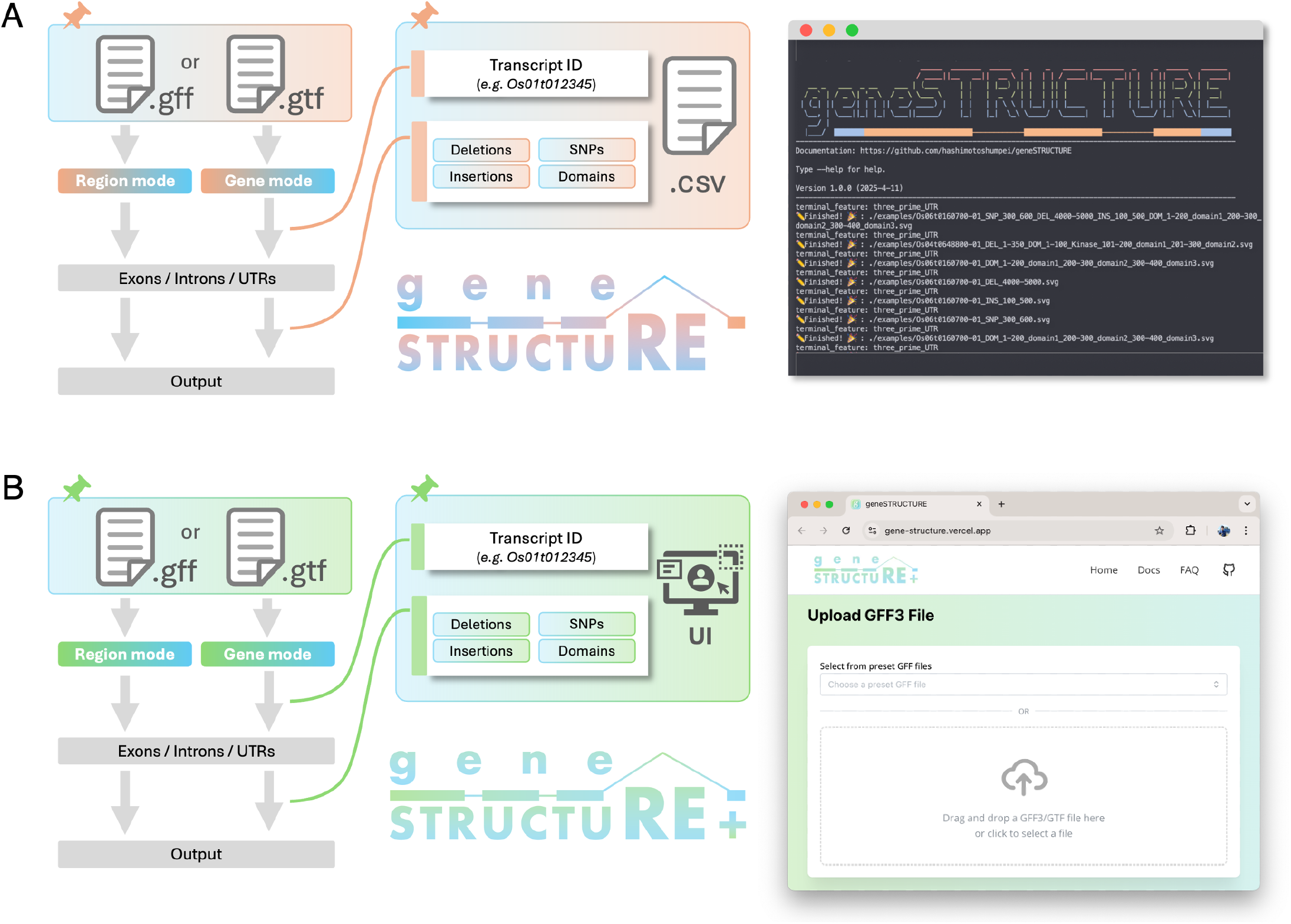
Workflows of geneSTRUCTURE (CLI) and geneSTRUCTURE+ (web application) Overview of the workflows implemented in geneSTRUCTURE (**A**) and geneSTRUCTURE+ (**B**). Both tools accept GFF3/GTF annotation files and support gene mode and region mode for visualizing exons, introns, and untranslated regions (UTRs). In the CLI version, transcript IDs and optional annotations, including insertions, deletions, SNPs, and protein domains, are provided via CSV files. In contrast, geneSTRUCTURE+ offers a browser-based user interface that enables interactive transcript selection, real-time visualization, and direct export of gene structure figures.

Fig. 2A presents an example of gene structure visualization generated by this tool. Here, the gene structure of *Os06g0160700* (*Os06t0160700-01*) is shown. Following conventional representation, exons (coding sequences, CDSs) and UTRs are depicted as rectangles, while introns are shown as connecting straight lines. In this tool, all transcripts are visualized in a unified orientation, with the 5′ end on the left and the 3′ end on the right, regardless of whether the gene is encoded on the plus or minus strand. To explicitly indicate this orientation, the terminal structure at the right end of the gene model is rendered with an arrow-like shape, thereby improving interpretability. In addition to basic gene structures, various user-defined annotation features can be overlaid onto the visualization. Deletions are represented as dashed chevron-shaped lines, insertions as downward-pointing arrows, and SNPs as vertical lines. Functional domains can also be incorporated and are displayed using user-specified colors. In addition to the “gene mode” described above, the tool provides a “region mode”. When this mode is enabled, all annotated transcripts located within a user-defined genomic interval are visualized simultaneously (Fig. 2B). Although additional gene-specific annotations cannot be added in this mode, the color schemes for structural elements such as exons and introns can be customized in the same manner as in gene mode.

**Fig 2.**
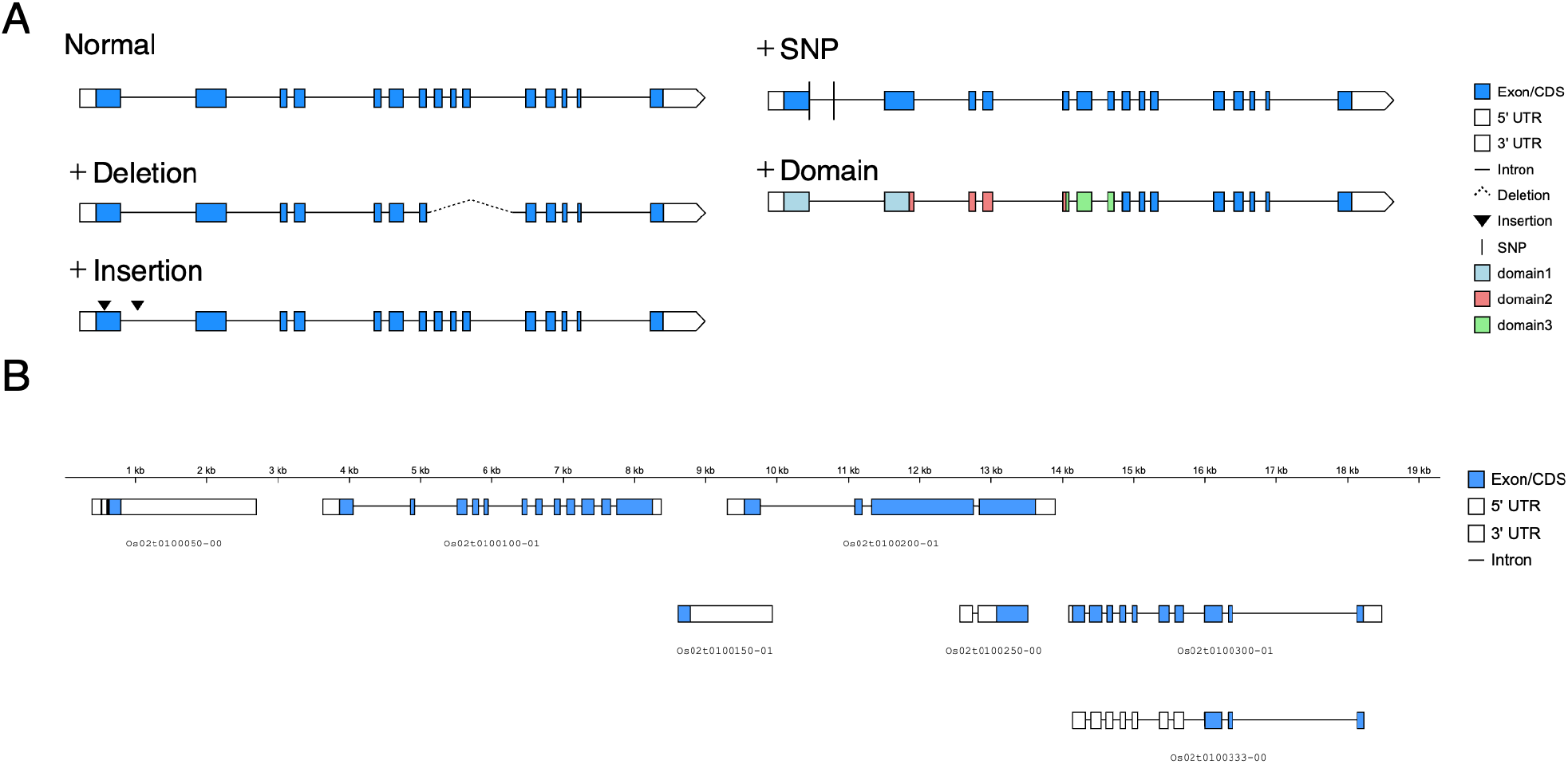
Gene structure visualizations generated by geneSTRUCTURE. **(A)** Gene mode visualizes the exon–intron organization of individual transcripts. It supports the overlay of additional annotations, including genetic variants (deletions, insertions, and SNPs) and protein domains, enabling detailed inspection of transcript-level structural and functional features. **(B)** Region mode displays multiple genes located within a user-defined genomic interval. Genes are aligned according to their genomic coordinates, facilitating comparison of gene organization and relative positions across a genomic region. In both modes, exons, introns, and untranslated regions (UTRs) are represented schematically.

Compared with conventional tools such as GSDS 2.0 (Hu et al., 2015), which operate exclusively as web servers, geneSTRUCTURE offers the advantage of local, standalone execution. Existing tools primarily focus on visualizing gene structures as described in annotation files. In contrast, geneSTRUCTURE extends this functionality by providing two major added values based on annotation data. First, it enables explicit visualization of genetic variants, including deletions, insertions, and SNPs. Second, it offers a region mode, in which genes within a user-defined genomic interval are displayed side by side along physical coordinates. The former function facilitates the organization and presentation of allelic variation, whereas the latter is particularly useful for visualizing candidate regions identified through QTL mapping or genome-wide association studies (GWAS). Through these added functionalities, geneSTRUCTURE is clearly distinguished from existing gene structure visualization tools.

The web-based interface is built with React, a widely adopted framework for building interactive user interfaces. React’s component-based architecture and optimized rendering mechanism enable efficient incremental screen updates, where only the affected visual elements are redrawn when users modify display parameters such as colors, spacing, or genomic regions. This unidirectional data flow ensures that state changes are consistently reflected across the interface without conflicts, which is essential for applications requiring high-frequency updates during real-time customization. Although React has become a dominant framework in general web development, its adoption in bioinformatics research tools remains relatively limited. A few notable examples in genome browsing, molecular visualization, and bacterial genome assembly have demonstrated how React enables smooth and responsive user experiences for complex biological data (Diesh et al., 2023; Grant et al., 2023; Sehnal et al., 2021). geneSTRUCTURE extends this approach to gene structure visualization, offering a modern, stable framework with an enhanced user experience.

## Acknowledgments

We thank Mr. Takumi Kawauchi (Laboratory of Plant Breeding and Genetics, University of Tokyo) for valuable comments on the manuscript. This research was partially supported by JSPS KAKENHI Grant Number JP23KJ0326.

## Author Contributions

S.H. conceived the project and developed the command-line interface version of the software. K.Y. developed the web-based application under the supervision of T.I. S.H. and K.Y. drafted the manuscript. S.H. revised and finalized the manuscript.

